# The fine-scale recombination rate variation and associations with genomic features in a butterfly

**DOI:** 10.1101/2022.11.02.514807

**Authors:** Aleix Palahí i Torres, Lars Höök, Karin Näsvall, Daria Shipilina, Christer Wiklund, Roger Vila, Peter Pruisscher, Niclas Backström

## Abstract

Genetic recombination is a key molecular mechanism that has profound implications on both micro- and macro-evolutionary processes. However, the determinants of recombination rate variation in holocentric organisms are poorly understood, in particular in Lepidoptera (moths and butterflies). The wood white butterfly (*Leptidea sinapis*) shows considerable intraspecific variation in chromosome numbers and is a suitable system for studying regional recombination rate variation and its potential molecular underpinnings. Here, we developed a large wholegenome resequencing data set from a population of wood whites to obtain high-resolution recombination maps using linkage disequilibrium information. The analyses revealed that larger chromosomes had a bimodal recombination landscape, potentially due to interference between simultaneous chiasmata. The recombination rate was significantly lower in subtelomeric regions, with exceptions associated with segregating chromosome rearrangements, showing that fissions and fusions can have considerable effects on the recombination landscape. There was no association between the inferred recombination rate and base composition, supporting a negligible influence of GC-biased gene conversion in butterflies. We found significant but variable associations between the recombination rate and the density of different classes of transposable elements (TEs), most notably a significant enrichment of SINEs in genomic regions with higher recombination rate. Finally, the analyses unveiled significant enrichment of genes involved in farnesyltranstransferase activity in recombination cold-spots, potentially indicating that expression of transferases can inhibit formation of chiasmata during meiotic division. Our results provide novel information about recombination rate variation in holocentric organisms and has particular implications for forthcoming research in population genetics, molecular/genome evolution and speciation.

## Introduction

The meiotic division process allows sexually reproducing organisms to generate haploid gametes. During meiosis, doublestrand breaks are induced into DNA, and a proportion (e.g. 5% in Arabidopsis thaliana, 10% in Mus musculus) of those are resolved as crossovers (Choi and Henderson 2015, Moens et al. 2002), leading to novel combinations of maternal and paternal chromosome segments. It is well established that the frequency and genomic distribution of crossovers can influence both micro- and macro-evolutionary processes – detailed knowledge about the recombination landscape is therefore key for understanding the relative importance of different proximate and ultimate mechanisms affecting genome evolution, generation and maintenance of genetic diversity, adaptation and speciation (Dapper and Payseur 2017, Stapley et al. 2017, Peñalba and Wolf 2020).

The frequency of recombination events resolved as crossovers (from here on recombination) can vary both at the inter- (Stapley et al. 2017, Smukowski and Noor 2011) and intra-specific level (Samuk et al. 2020), as well as between individuals within populations (Johnston et al. 2016) and within individuals over time (Stapley et al. 2017, Peñalba and Wolf 2020). It is well established that the recombination rate has a genetic component, but the rate can also be influenced by environmental factors (i.e., recombination is partly plastic; Peñalba and Wolf 2020). Recombination rate can also vary considerably across chromosomes and chromosome regions in many species, and mapping this variation may shed light on the mechanistic control of where in the genome recombination is initiated. The reasons for such regional recombination rate variation have been studied in detail in a few organism groups and a handful of consensuses have been reached. First, the size of a chromosome can affect the recombination rate, mainly because correct segregation seems to be dependent on at least one recombination event per chromosome arm in many species (Pardo-Manuel de Villena and Sapienza 2001, Smith and Nambiar 2020). A lower recombination rate on the sex-chromosomes than on the autosomes is also a commonly observed pattern and this difference can potentially be attributed to low sequence homology as a consequence of general degeneration of sex-limited chromosomes (Y or W) (e.g. Bergero and Charlesworth 2009). Within chromosomes, the location of recombination events might be determined by preferential initiation of double-strand breaks close to the telomeres and where the chromatin in general is open and accessible (e.g. Haenel et al. 2018, Gray and Cohen 2016). Conversely, the recombination rate is usually suppressed within and around the highly heterochromatic centromeres (Dapper and Payseur 2017, Stapley et al. 2017). Furthermore, physical interference between multiple chiasmata may lead to regional differences in the recombination rate (Gray and Cohen 2016, Peñalba and Wolf 2020). In mammals, the gene PRDM9 mediates recombination by binding to specific sequence motifs, which explains that most recombination events occur in a limited portion of the genome, i.e. recombination hot-spots (e.g. Myers et al. 2005, Grey et al. 2011). However, most vertebrate and all evertebrate lineages lack a functional copy of PRDM9 and recombination initiation must hence be mediated by other factors in these species.

We expect the regional variation in recombination rate to be associated with genomic features, the efficiency of selection and the levels of genetic diversity (Dolgin and Charlesworth 2003, Petrov et al. 2011). Such regional variation in the efficiency of selection can for example affect the distribution of transposable elements (TEs), which tend to accumulate in regions of low recombination rate in both animals (Bartolomé et al. 2002) and plants (Xu and Du 2014). In addition, the frequency of recombination events can also affect nucleotide composition as a direct consequence of GC-biased gene conversion (gBGC), a process that facilitates the fixation of strong (G and C) over weak nucleotides (A and T) during the double-strand break repair step (Duret and Galtier 2009). Although genome-wide estimates of the recombination rate and large-scale variation landscapes have been obtained for many species, detailed recombination maps are still mainly limited to model organisms and domesticated species, and little is known about recombination rate variation in natural populations. Hence, a broader taxonomic sampling will be needed to get a more complete picture of what drives recombination rate variation within genomes and between lineages. This applies not the least to holocentric organisms which have chromosomes without localized centromeres (Suomalainen 1953), such as Lepidoptera (butterflies and moths), where the research on causes and consequences of recombination rate variation is still in its infancy.

Here, we used a large whole-genome re-sequencing data set from a Swedish population of the wood white butterfly (*Leptidea sinapis*) to characterize the fine-scale variation in recombination rate and assess potential associations between recombination rate, nucleotide composition and genomic features. So far, the knowledge about regional recombination rate variation in Lepidoptera is restricted to a handful of pedigree based genetic maps (Davey et al. 2017, Shipilina et al. 2022, Smolander et al. 2022) and this is therefore a spearheading attempt to describe the fine-scale variation in recombination rate and the potential associations with genomic features in a butterfly. The wood white is widely distributed across western Eurasia and shows extreme intraspecific variation in chromosome numbers, with an increasing number of chromosomes in a cline-like pattern from 2n ~ 56 – 60 in the northern (Scandinavia) and eastern (central Asia) parts of the distribution range to 2n ~ 106 – 110 in the south-western (Iberia) part of the distribution range (Dincă et al. 2011, Lukhtanov et al. 2018). Hence, some wood white populations differ significantly from the ancestral Lepidoptera chromosome number of n = 31 (Robinson 1971, Ahola et al. 2014). Previous studies suggest that recurrent chromosome fissions and fusions underlie this variation (Dincă et al. 2011, Talla et al. 2017) and that at least a handful of fission/fusion polymorphisms segregate in the Swedish population (Höök et al. 2022). The rapid karyotype evolution in wood whites provides a unique system for characterizing the regional variation in recombination rate in a natural population of a holocentric species, and combining it with investigating the effects of segregating chromosome rearrangements on the recombination landscape. Given the key role of recombination in both micro- and macro-evolutionary processes, our results will be important for understanding molecular mechanisms and evolutionary forces affecting genome evolution and divergence processes in holocentric organisms in general.

## Results

### Demographic history inferences

The demographic trajectories inferred separately for each chromosome jointly revealed the demographic history of the Swedish population of *Leptidea sinapis.* These demographic trajectories shared three main features. Firstly, a maximum effective population size (*N_e_*) of around 106 approximately 10,000 generations before present (BP), preceded by a period of high *N_e_* and a slight increase matching with the end of the Last Glacial Period. After this time point, *N_e_* started to decline exponentially until stabilizing about 100 generations BP. In the most recent past, *N_e_* remained constant. Contemporary estimates of *N_e_* oscillate between 10^3^ and 2 * 10^4^ for the different chromosomes (Supplementary Figure 1).

### Recombination rate variation and distribution

The estimated genome-wide recombination rate was 7.37 cM/Mb, with measurements for individual chromosomes ranging between 3.5 - 15.3 cM/Mb. There was a marginally nonsignificant (Spearman *ρ* = −0.292, *p* = 0.06) negative association between the recombination rate and chromosome length (Supplementary Figure 2). Autosomes (7.65 cM/Mb) showed on average a higher recombination rate than the Z-chromosomes (7.03 cM/Mb), although this difference was not significant (Wilcoxon’s test, W = 41, p = 0.92).

Visual inspection of the variation in recombination rate revealed a considerably reduced recombination rate towards chromosome ends (Figure 1A). A formal analysis showed that the recombination rate in the subtelomeres (last 5 100 kb windows at each end of the chromosomes) was significantly reduced (2.46 cM / Mb) compared to the 100 kb windows located in proximal positions of the chromosomes i.e., outside of the subtelomeric regions (Wilcoxon’s test, W = 1,310,856, p-value = 3.18 * 10^−66^). It should be noted that this does not necessarily reflect a low recombination rate in the telomeres, but rather in the subtelomeric regions, since telomeric repeats were not covered in the wood white genome assembly (Höök et al. 2022). We also analyzed each chromosome separately and found a reduction in recombination rate in subtelomeric regions in 50 out of the 58 chromosome ends (29 chromosome pairs, 2n = 58; Supplementary Table 1). Out of the 8 chromosome ends that did not show a reduced recombination rate, four match observations of segregating fission/fusion polymorphisms – these involve chromosome pairs 18 and 25, 11 and 26, and 5 and 27, respectively (Höök et al. 2022).

**Figure 1.**
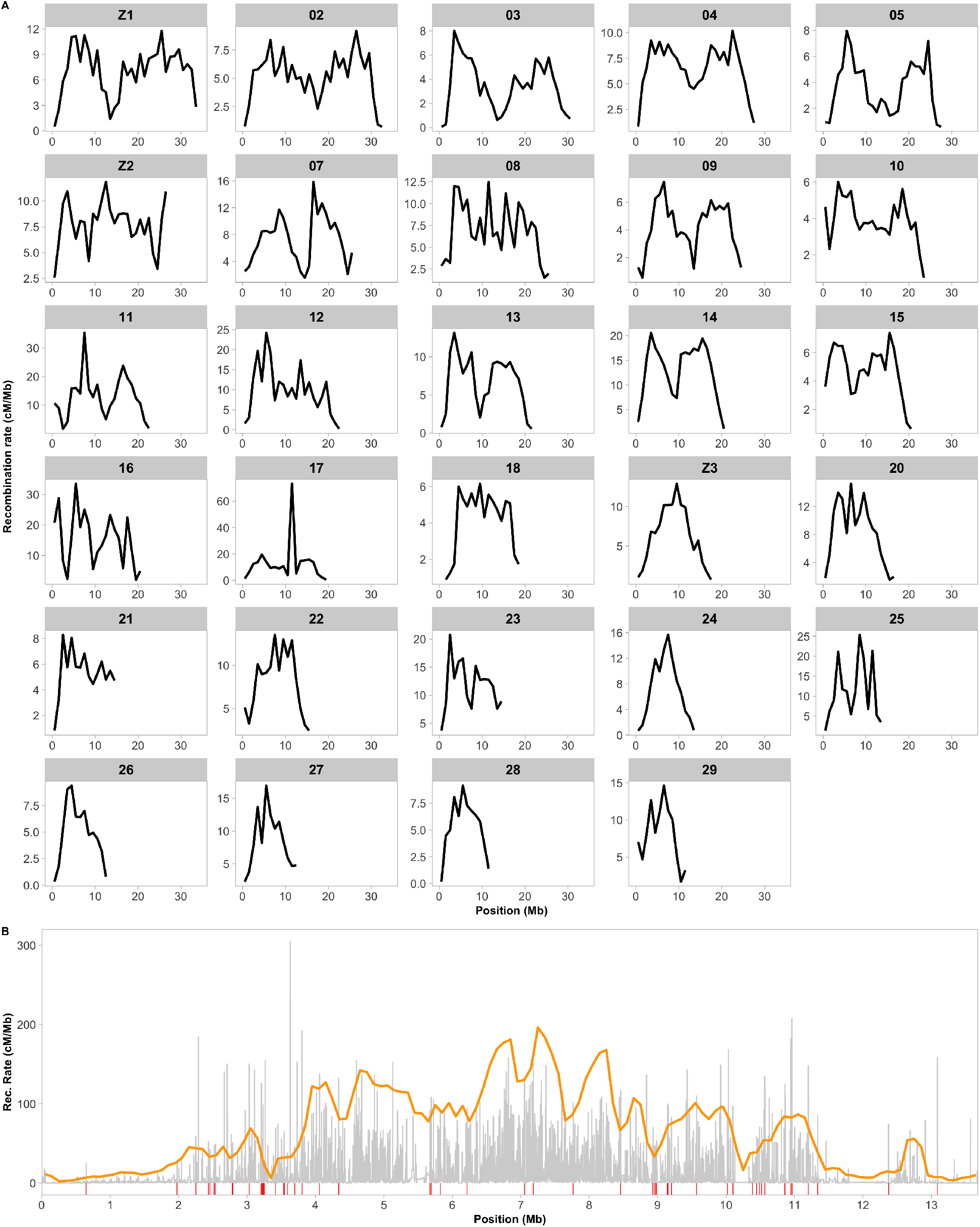
Recombination rate estimations and hot-spot determination. (A) The 1Mb-scale estimates of recombination rate in 1Mb windows are shown in cM/Mb for all chromosomes, ordered by decreasing length. (B) Detailed recombination map for chromosome 24, with all oscillations in recombination rate inferred by pyrho in grey. The orange line represents the 10x background recombination rate, the threshold used to determine the minimum recombination rate of local hot-spots. Red lines underneath the plot indicate the presence of a recombination hot-spot according to our defining parameters. See Materials & Methods for a more detailed description of the parameters.

Besides a significantly reduced recombination rate in subtelomeric regions in almost all chromosomes, we also found that regional variation differed in other respects. First, some chromosomes showed an obvious unimodal distribution for the recombination rate, with a maximum value in the central region and a progressive decrease towards the terminal regions. This was a frequent (but not exclusive) pattern for the shorter chromosomes (Figure 1A). In contrast, the recombination rate was bimodally distributed along some chromosomes, with a central region of reduced recombination in addition to the reduction in subtelomeric regions. This was the most commonly observed pattern for the larger chromosomes (Figure 1).

### Recombination hot-spot and cold-spot identification

Informed by coalescent simulations, we developed thresholds for identification recombination hot- and cold spots, which take into account both specific demographic history of the population of interest and the potential stochasticity of our method of recombination rate inference (see methods for further details). Based on these thresholds, a total of 3,124 recombination hotspots were classified (Figure 1B). The hot-spots had an average length of 1,656 bp and the mean recombination rate within hotspots was 94.1± 62.5 cM/Mb. The highest estimated rate in any hotspot was 708 cM/Mb. The average recombination rate for hot-spots represented an approximate 13-fold increase over the genome-wide recombination rate, but hot-spots only constituted 5.2 Mb (0.87%) of the genome. Recombination hot-spots were found at a significantly lower frequency in the terminal 10% regions of the chromosomes (5% on each end); Wilcoxon’s W = 220.5; p = 4.94 * 10^-4^ (Supplementary Figure 3A). A higher density of recombination hot-spots was detected on the three Z-chromosomes (six hot-spots / Mb) when compared to the autosomes (five hot-spots / Mb), but the difference was not significant (Student’s t-test, t = −0.85556, df = 27, p = 0.20. Permutation analysis revealed a significantly lower LINE (p = 0.02) and LTR density (p > 0.001) and a higher SINE density (p < 0.001) in the hot-spots, while DNA transposon density did not deviate significantly (p = 0.49) from the genome wide average (Figure 2). There was no enrichment of functional gene categories or specific sequence motifs in the hot-spots (p < 0.05, Supplementary Table 2).

**Figure 2.**
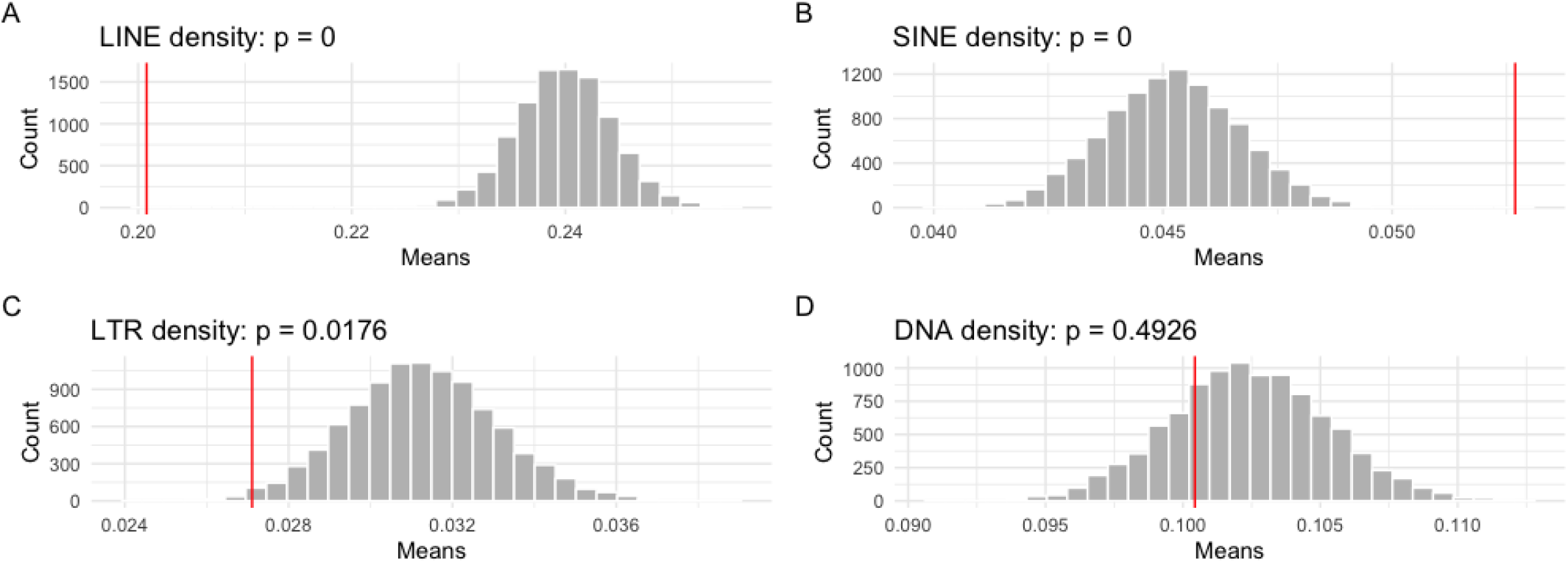
Density of TEs in hot-spots. The bars show the distribution of sampled TE density means for 3,124 random 1,656 bp genomic windows (number and average length of the hot-spots). The red line indicates the observed mean TE densities in the defined hot-spot windows.

We also identified 1,283 recombination cold-spots, i.e., regions with considerably reduced recombination rate as compared to the genomic average (see methods). The average length of the cold-spots was 30 kb, and 70 of the cold-spots were longer than 100 kb. Despite the lower frequency of cold-spots than hot-spots, they represented a substantially larger proportion of the genome (38.2 Mb, 6.37%). As expected given the overall reduced recombination rate in subtelomeric regions, cold-spots were particularly abundant in these regions. In particular, subtelomeric regions contained 29.6% of the cold-spots in total, while representing only 4.8% of the genome. The cold-spots located outside subtelomeric regions had an average length of 25.6 kb and 46 were longer than 100 kb. This translates to a significant enrichment in the frequency of cold-spots in the 10% most terminal chromosomal regions compared to the more central positions of the chromosome (Wilcoxon’s W = 3; p = 5.62 * 10^-4^) (Supplementary Figure 3B). Similar to the observation for hotspots, there was no significant difference in cold-spot frequency between the autosomes (1.8 cold-spots / Mb) and the three Z-chromosomes (2.5 cold-spots / Mb) (Student’s t-test, t = −0.74483, df = 27, p-value = 0.231. Permutation analyses revealed a significantly lower transposon density in cold-spots, consistent across all classes (p < 0.01; Figure 3). There was also a significant enrichment of genes related to transtransferase activity in the cold-spot regions compared to the genome in general (Supplementary Table 3), but no enrichment of specific sequence motifs in the cold-spots (p > 0.05).

**Figure 3.**
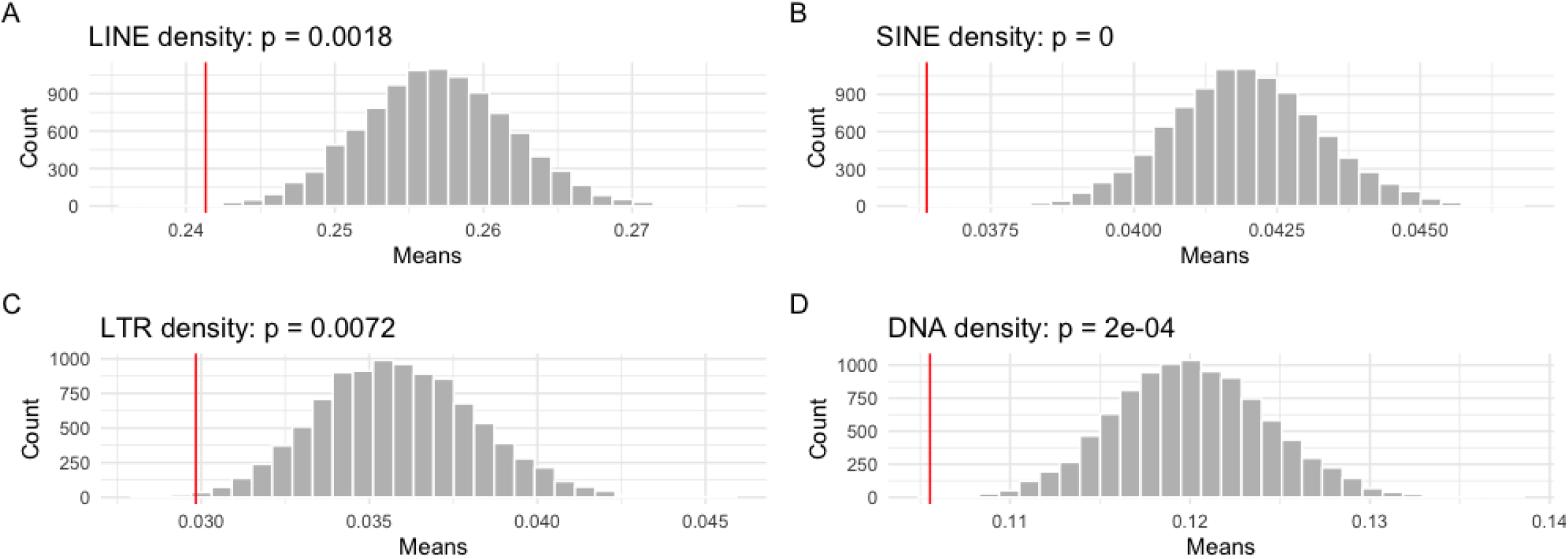
Density of TEs in cold-spots. Histogram shows the distribution of permuted TE density means for 1,283 random 30 kb genomic windows (number and average length of the cold-spots). The red lines indicate the the observed mean TE densities in the defined cold-spot windows.

### Association between recombination rate and base composition

To investigate potential effects of gBGC on the base composition in the wood white genome, we assessed the relationship between the recombination rate and nucleotide composition using a window-based approach. The analysis showed that the recombination rate (averaged over 1 Mb windows) was not significantly correlated with the GC content in the genome in general (Spearman *ρ* = −0.07, p = 0.10) (Figure 4A).

**Figure 4.**
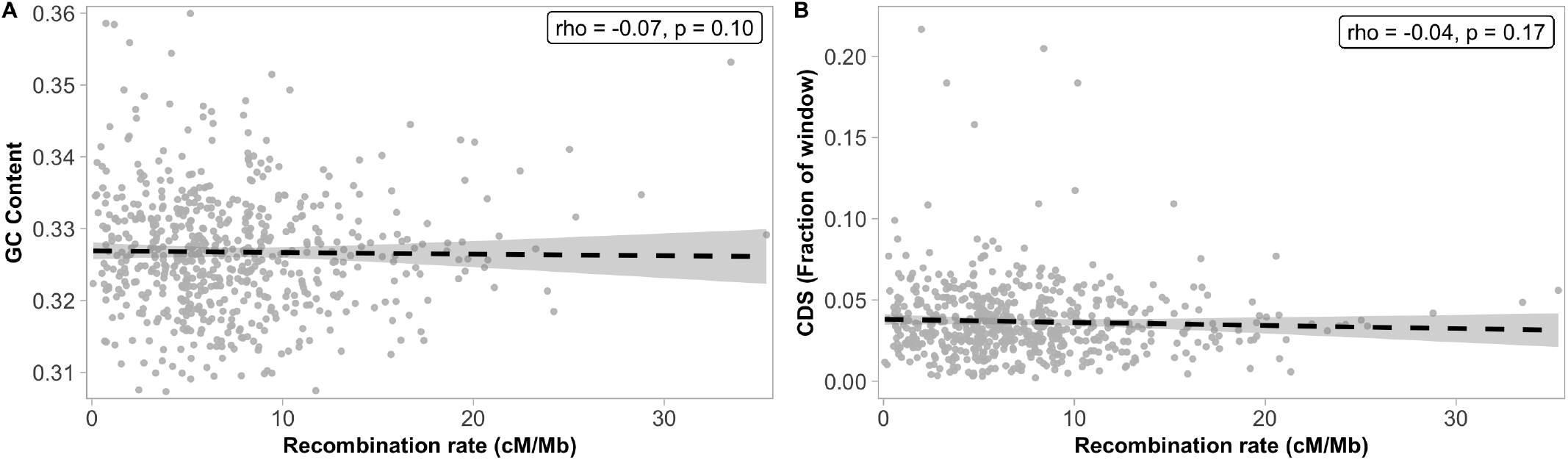
Associations between the recombination rate and base composition (A) and gene density (B). Correlation between parameters was calculated using 1 Mb genomic windows.

Since we observed a significant reduction in recombination rate in subtelomeric regions, we investigated if those regions had deviating base composition. The analysis showed that there was a significantly higher GC content in subtelomeric regions (33.46%) compared to proximal chromosome regions (32.61%) (Wilcoxon’s W = 1,109,810; p = 5.73 * 10^-22^).

The GC content was also slightly lower within hot-spots (32.43%) and their 5 kb flanking regions (32.45%) as compared to the genome-wide estimates (32.65%).

### Associations between recombination rate and genomic features

We used multiple regression to investigate the relative effect of different explanatory variables (GC content, gene density, DNA transposon, SINE, LINE and LTR retrotransposon densities). The regression model revealed an overall significant association between recombination rate and the explanatory variables (F(6) = 10.77, df = 603, p = 2.13 * 10^-11^), but it explained a marginal part of the total variation in the recombination rate (R^2^ = 0.10, *AdjR*^2^ = 0.09) and only SINE and LINE density were significant explanatory variables (Table 1).

**Table 1.** Linear regression model with six explanatory variables included. Explanatory variables that are significantly associated with recombination rate variation at a 1 Mb scale are highlighted in bold.

|  | Estimate | Std. error | t | p-value |
| --- | --- | --- | --- | --- |
| GC | 27.5203 | 24.6317 | 1.117 | 0.264 |
| Gene density | -6.6062 | 6.7692 | -0.976 | 0.329 |
| DNA | 7.2352 | 16.7373 | 0.432 | 0.666 |
| **LINE** | -30.4439 | 7.2939 | -4.174 | $3.44e^{-05}$ |
| **SINE** | 104.8162 | 16.0648 | 6.525 | $1.45e^{-10}$ |
| LTR | 32.1472 | 23.9983 | 1.340 | 0.181 |

The recombination rate was not significantly associated with genome-wide gene density (Spearman *ρ* = −0.04, p = 0.17; Figure 4B). However, when partitioning the data and running analyses for different gene elements separately, we observed a significantly lower (Wilcoxon’s W = 1.5971*e*^-10^, p-value < 2.2*10^-16^) recombination rate within exons (5.5 cM/Mb) and in 5’ UTR regions (6.3 cM/Mb; Wilcoxon’s W = 1.5869 * 10^8^, p-value < 2.2*10^-16^) and a significantly higher recombination rate in introns (Wilcoxon’s W = 7.3462*10^12^, p-value < 2.2*10^-16^) compared to intergenic regions (7.5 cM/Mb) (Figure 5).

**Figure 5.**
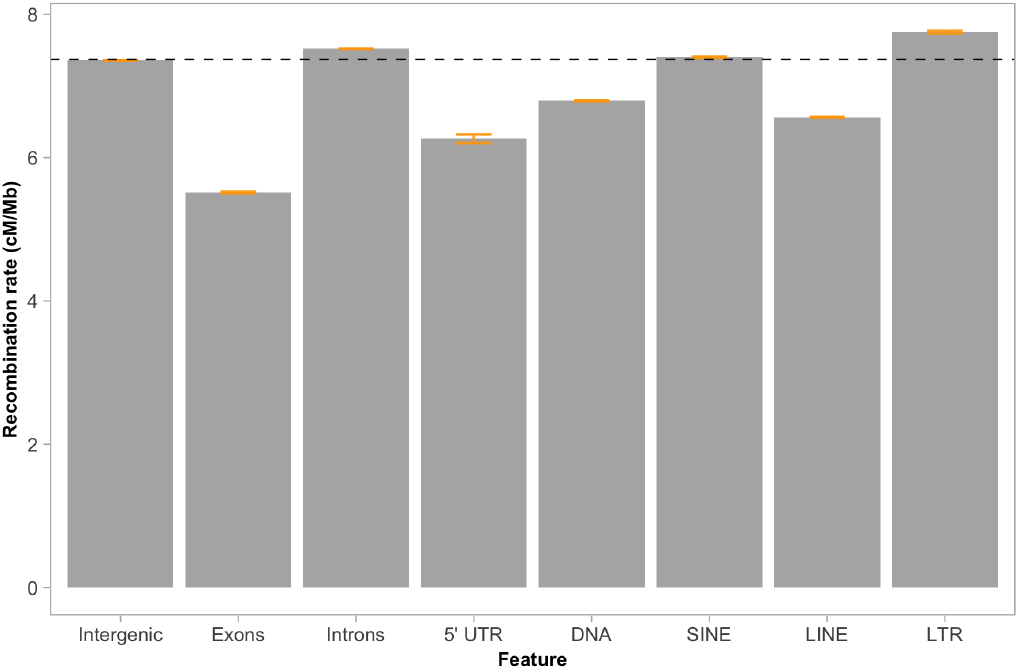
Recombination rate estimates in different gene elements and classes of transposable elements. The genome-wide recombination rate (7.37 cM/Mb) is indicated with the horizontal black dashed line. Orange bars indicate the 95% confidence intervals.

The associations between recombination rate and the densities of different TE classes varied considerably. For all four classes analyzed, we found a significant correlation with the recombination rate, but the direction and strength of these associations varied. DNA transposons (Spearman *ρ* = 0.09, p = 0.03) and SINEs (Spearman *ρ* = 0.29, p = 3.31*10^-13^) were positively associated, and LTRs (Spearman *ρ* = −0.11, p = 8.50*10^-3^) and LINEs (Spearman *ρ* = −0.19, p = 3.01 *10^-6^) negatively associated with the recombination rate (Figure 6).

**Figure 6.**
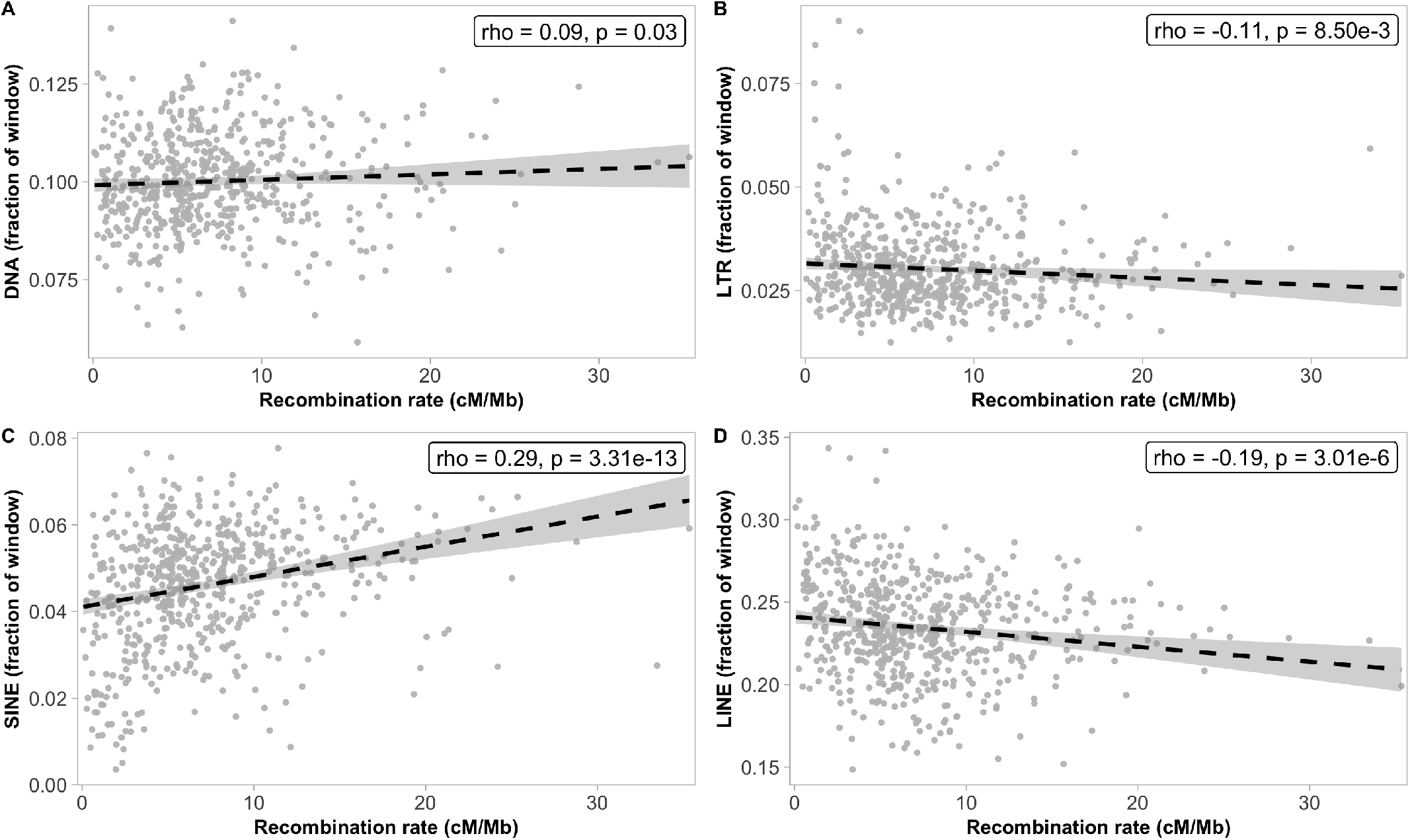
Association between the recombination rate (1 Mb scale) and the proportion of four different TE classes; (A) DNA transposons, (B) LTRs, (C) SINEs and (D) LINEs. The abundance of TEs was calculated as the fraction of each 1 Mb window occupied by each specific element.

Differences between classes of transposable elements were not restricted to the overall associations with recombination rate. The average recombination rate within each class of TEs varied as well. In LINEs (6.6 cM / Mb; Wilcoxon’s W = 5.233*1012, p-value < 2.2*10-16) and DNA transposons (6.8 cM / Mb; Wilcoxon’s W = 2.6104*1012, p-value < 2.2*10-16) the recombination rate was significantly lower, and in SINEs (7.4 cM / Mb; Wilcoxon’s W = 1.2212*1012, p-value < 2.2*10-16) and LTRs (7.8 cM / Mb; Wilcoxon’s W = 7.0468*1011, p-value < 2.2*10-16) the recombination rate was significantly higher than the genomic average rate (Figure 5).

## Discussion

### Genome-wide distribution of the recombination rate in wood whites

The genome-wide rate of recombination has been shown to vary considerably between different insect species, from comparatively low in Diptera (< 1 cM / Mb, Beye et al. 2006) to exceptionally high in honeybees (19 cM / Mb; Beye et al. 2006). We found that the genome-wide recombination rate in the Swedish wood white population was 7.37 cM / Mb. This is slightly higher than estimates from other lepidopterans like *Bombyx mori* (4.6 cM / Mb), *Heliconius melpomene* (5.5 cM / Mb) and *H. erato* (6 cM / Mb) (Tobler et al. 2005; Yasukochi 1998; Jiggins et al. 2005) and substantially higher than in vertebrates for which estimates range between 0.16 cM / Mb in the Atlantic trout to 3.17 cM / Mb in chicken (Beye et al. 2006).

Although not significant at the 5% level, we found that the recombination rate was negatively associated with chromosome size. A clearly deviating rate was found for chromosome 16 (15.3 cM / Mb), which explains why the association between chromosome size and recombination was non-significant. Negative associations between recombination rate and chromosome length have repeatedly been observed in different taxonomic groups, for example yeast (Kaback et al. 1992), humans and rodents (Jensen-Seaman et al. 2004), birds (Backström et al. 2010), cattle (Mouresan et al. 2019) and butterflies (Martin et al. 2019, Shipilina et al. 2022). Such a relationship is expected given that crossovers are necessary for correct segregation of chromosomes during meiosis (Pardo-Manuel de Villena and Sapienza 2001, Smith and Nambiar 2020). Our analysis also showed that longer chromosomes tended to have a bimodal recombination rate distribution with a reduced rate at the chromosome center and towards the chromosome ends. This pattern is in line with findings in other taxa, both organisms with defined centromeres where recombination is reduced (Dapper and Payseur 2017), and holocentric species such as *Caenorhabditis elegans,* in which the recombination rate has been shown to increase with the relative distance from the center of the chromosomes (Prachumwat et al. 2004). The reduced recombination rate in the center of chromosomes in monocentric species is often a direct consequence of the lack of crossing-over events in the centromeres. This explanation is obviously not valid in holocentric lineages. Since the pattern is restricted to larger chromosomes, a potential explanation to the reduced rate in chromosome centers could be the occurrence of multiple chiasmata on larger chromosomes. For example, observations in *Psylla foersteri* suggest that longer chromosomes can accommodate the formation of two simultaneous chiasmata, while shorter chromosomes only have one (Nokkala et al. 2004). In the cases where two chiasmata are formed in a single chromosome, crossover interference may prevent those from forming near each other and tend to drive them towards opposite ends of the chromosome (Otto and Payseur 2019). An additional, but not mutually exclusive explanation, is the formation of the “meiotic bouquet”, a stage in early meiosis characterized by the aggregation of chromosome ends close to the nuclear membrane, which can drive the crossovers towards distal positions and reduce the recombination rate in the center of the larger chromosomes (Scherthan et al. 1996, Haenel et al. 2018).

Besides a reduced recombination rate in the center of larger chromosomes, we also found a reduced recombination rate towards the very ends of chromosomes. This pattern was observed for almost all chromosomes (see exceptions below), irrespective of chromosome size and type. Note that the telomeres, which in Lepidoptera are 6-8 kb long tandem repeats of the motif (TTAGG)n (Okazaki et al. 1993, Sasaki and Fujiwara 2000) were not assembled in the reference genome. Such decrease in the recombination rate in the subtelomeric regions of the chromosomes has previously been observed in Heliconius butterflies (Martin et al. 2019), and also in some other organism groups such as yeast (DuBois et al. 2002, Barton et al. 2008) and flycatchers (Kawakami et al. 2013). A potential explanation for this pattern is that crossover initiation is prevented near the telomeres to minimize the risk for ectopic recombination between non-homologous repeat sequences during meiosis (Smith and Nambiar 2020). As mentioned above, our results showed that the reduced recombination rate towards chromosome ends was not ubiquitous across all chromosomes; eight chromosome ends (one for each of the following chromosomes: 5, 6, 10, 11, 16, 25, 27 and 29) did not show a significant reduction in the recombination rate as compared to each respective intra-chromosomal level. Four of these exceptions (one chromosome end for each of chromosomes 5, 11, 25 and 27) coincide with recently identified fission and fusion polymorphisms segregating in the wood white population in Scandinavia (Höök et al. 2022). These results show that fission/fusion events can have immediate effects on the distribution of crossover events within and between chromosomes.

### Characterization of recombination hot-spots and coldspots

The total number of hot-spots (n = 3,124) identified in the wood whites is equivalent to what has previously been observed in for example Ficedula flycatchers using a comparable approach (Kawakami et al. 2013). The density of hot-spots was, however, much lower than in humans (n = 25,000 - 50,000 hot-spots in an approximately five times larger genome) (Myers et al. 2005). Although the specific thresholds for defining hot-spots vary between studies, they all rely on the comparison of the background and local recombination rates, making the results reasonably comparable. We found that the distribution of hot-spots was similar between wood whites and humans, as hot-spots occurred mostly outside of genes (McVean et al. 2004). This is in contrast to birds, which show an enrichment of hot-spots within genic and regulatory regions (Kawakami et al. 2013; Singhal et al. 2015; Smeds et al. 2016). We found that the frequency of recombination hot-spots and cold-spots was relatively similar in the center of chromosomes, but that the number of hot-spots decreased, and cold-spots were more frequent towards chromosome ends. A higher occurrence of recombination cold-spots in terminal regions of the chromosomes has previously been observed in yeast, which seem to lack crossovers close to chromosome ends altogether (Su et al. 2000; Barton et al. 2003). This observation is also in line with the observed decrease in average recombination rate close to chromosome ends in the wood whites and again suggests that the recombination machinery is partly blocked from accessing the very ends of chromosomes.

We found that the recombination landscape in the wood white was highly variable. This is in line with observations in other organisms like humans, birds (Singhal et al. 2015) and dogs (Axelsson et al. 2012). However, other insects like for example *D. melanogaster* (Comeron et al. 2012), for which detailed recombination maps are available, generally show less pronounced recombination hot-spots. While the hot-spot locations in humans largely are determined by the presence of sequence motifs associated with PRDM9 binding (Grey et al. 2011), little is known about what drives crossovers to occur at specific locations in organisms that lack a functional copy of PRDM9. In order to get preliminary information about potential mechanistic underpinnings of recombination rate variation in wood whites, we therefore assessed if specific sequence motifs or gene categories were enriched in recombination hot-spots and cold-spots. The analyses did not reveal any associations for sequence motifs, and we found no enrichment of specific gene categories in hot-spots. In cold-spots, however, there was a significant enrichment of genes with functions associated to transferase activity. Particularly interesting is the case of farnesyltranstransferase activity, as farnesylation is a key step for the correct attachment between the spindle and the kinetochores in humans (Moudgil et al. 2015). It is therefore tempting to speculate that the active expression of farnesyltranstransferase might block the recombination machinery close to those genes. However, since farnesyltranstransferases are located in only 13 different cold-spot regions, other forces must also underlie the absence of recombination in many cold-spots.

### Associations between recombination, nucleotide composition and gene content

The high-density recombination maps developed here, allowed us to investigate potential associations between the local recombination rate and different genomic features. Such information can be used to deduce the effects of recombination on base composition and/or potential regulatory mechanisms modulating the recombination landscape. A potential driver of a positive association between recombination and nucleotide composition is GC-biased gene conversion (gBGC), i.e., the fixation bias favoring “strong” alleles (G and C) over “weak” alleles (A and T) during meiotic recombination (Duret and Galtier 2009). This process mimics directional selection and can lead to deviating nucleotide composition between regions experiencing different recombination rates. The analysis showed that the local recombination rate was not associated with nucleotide composition (GC-content) in the wood whites. We also found that recombination hot-spots had marginally lower GC-content (the opposite is usually observed when biased fixation is a considerable force; Kawakami et al. 2013). This is in line with a limited effect of gBGC in Leptidea butterflies (Boman et al. 2021) and stays in contrast to findings in several other systems like humans (Fullerton et al. 2001, Meunier and Duret 2004), mice (Clément and Arndt 2013), flycatchers (Kawakami et al. 2013) and fruit flies (Marais et al. 2001), as well as plants (Muyle et al. 2011) and yeast (Gerton et al. 2000, Kiktev et al. 2018). As far as we are aware, there are no other studies that have analyzed the strength of gBGC in butterflies outside the Leptidea genus. Hence, it is premature to draw conclusions regarding the impact of gBGC on nucleotide composition in Lepidoptera in general.

In many organisms, for example mouse (Paigen et al. 2008) and different plant species (Gaut et al. 2007, Tiley Burleigh 2015), recombination occurs more frequently in gene-dense genomic regions, but we did not find such an association in the wood whites. However, our data showed that the recombination rate was significantly reduced in exons and 5’ UTR regions compared to the introns and intergenic regions. This is in line with findings in other insects (Wallberg et al. 2015; Jones et al. 2019), as well as in humans (McVean et al. 2004), where recombination hot-spots mainly occur in the vicinity of, but not within, coding and regulatory regions. The small but significantly elevated recombination rate in introns compared to intergenic regions is consistent with findings in the holocentric nematode *C. elegans*(Prachumwat et al. 2004), but in contrast to the observations in for example Drosophila (Carvalho Clark 1999) and humans (Comeron Kreitman 2000). Taken together, this indicates that recombination occurs within genes in butterflies, but that crossovers are partly inhibited in coding sequences which might lead to a slightly elevated rate in introns.

### Different TE classes show contrasting association patterns with the recombination rate

Potential associations between recombination and TE densities have mainly been investigated in organisms with defined centromeres (Kent et al. 2017), while investigations in holocentric species are scarce (but see Lavoie et al. 2013, Baril Hayward 2022, Smolander et al. 2022). To investigate the potential associations between the recombination rate and genomic features in the wood white, we used density information for TEs previously identified in the species (Höök et al. 2022). The analysis revealed that associations between the recombination rate and the abundance of TEs varied considerably depending on the TE class. DNA transposons and SINEs were positively associated, while LINEs and LTRs were negatively associated with the recombination rate. Under the assumption that TE insertions in general are slightly deleterious we would expect a negative correlation between the recombination rate and the abundance of TEs (Kent et al. 2017), as a consequence of more efficient purging of deleterious insertions in regions with a higher recombination rate (Bartolomé et al. 2002, Wright et al. 2003). However, given that recombination is initiated by a double-strand break, it is possible that certain types of TEs are used as a template for the repairing process, driving them to higher frequencies in regions of high recombination rate (Onozawa et al. 2014). SINEs have for example have been shown to use DNA breaks to integrate back into the genome after replication (Singer 1982). Potential associations between TEs and the recombination rate are hence expected to depend on the occurrence of specific classes of TE in the focal study system. For example, Alu elements (a subfamily of SINEs) in humans have been shown to accumulate in regions with elevated recombination (Witherspoon et al. 2009) and SINEs are strongly positively associated with the recombination rate in the painted lady (*Vanessa cardui*) (Shipilina et al. 2022). Similarly, DNA transposons are associated with high recombination rate in C. elegans (Duret et al. 2000). The causality of such associations between the variation in recombination rate and the abundance of TEs is not easy to establish. In cases where TE proliferation has deleterious fitness effects, we expect a negative association between TE abundance and recombination rate. However, presence of Alu elements in humans has been shown to lead to an increase in the local recombination rate, possibly a consequence of that the Alu elements mimic the action of short recombinogenic motifs (Witherspoon et al. 2009). It is also possible that other underlying factors affect the TE and recombination rate distributions similarly – both the TE proliferation and the recombination initiation machinery for example seem to target open chromatin more easily (Kawakami et al. 2013). In the wood whites, the average recombination rate estimates within TE classes did not deviate considerably from the genome-wide average. However, these comparatively minor differences in the recombination rate can indicate differences in the selective pressure against insertion of specific families of TEs. This does not seem to be related with the length of the TEs, as SINEs – which are considerably shorter than LINEs and LTR elements – showed an average recombination rate between the longer types.

In summary, the different TEs showed different associations with the local recombination rate. These results are consistent with findings in other studies and may point toward similar determinants in holocentric organisms compared to those with defined centromeres. However, the causality needs further study, for example by detailed characterization of cross-over regions in large pedigrees.

## Materials and methods

### Genome assembly

The wood white genome assembly used as reference was developed for another study (Höök et al. 2022). In brief, one mated adult female wood white was caught in Sweden and kept in the lab for egg laying. From the offspring, one male pupa was sampled and flash frozen in liquid nitrogen. The sample was divided to create a 10X Genomics Chromium Genome-library and a Dovetail HiC-library from the same individual. For 10X sequencing, DNA was extracted using a modified HMW salt extraction method (Aljanabi and Martinez 1997). Tissue for HiC-sequencing was disrupted in liquid nitrogen. Library preparations, sequencing and genome assembly was performed by NGI Stockholm. Sequencing was performed on Illumina NovaSeq6000 with a 2×151 setup. 10X linked reads were assembled with 10X Genomics Supernova v2.1.0 (Weisenfeld et al. 2017). HiC reads were processed with Juicer v1.6 (Durand et al. 2016a) and used for scaffolding the 10X assembly with 3DDNA v.180922 (Dudchenko et al. 2017). Resulting assemblies were reviewed with Juicebox v1.11.08 (Durand et al. 2016b). In addition to minor corrections to the initial assembly, two chromosome sized scaffolds were merged. The assembly was finalized with the script ‘run-asm-pipeline-post-review.sh’ from the 3DDNA pipeline v.180922 (Dudchenko et al. 2017).

### Gene annotation

Gene annotation lift-over was performed by aligning wood white protein queries generated by Talla et at.(2017) to the current version of reference genome, using spaln 2.4.0 (Iwata and Ghoto 2012) with the parameters -Q7 -LS -O7 -S3.

### TE annotation

Repetitive element consensus sequences were predicted *de novo* using RepeatModeler 1.0.11 (Bao et al. 2015). Transposable elements characterized as unknown were submitted to CENSOR (Bao et al. 2015) for annotation, where any hits with a score < 200 were removed. All predicted sequences were matched against gene annotations using diamond blast 2.0.4 (Buchfink et al. 2021), to correct for annotation errors (bitscore > 100). Transposable elements were then annotated in the genome with RepeatMasker 4.1.0, using the predicted library of consensus sequences in wood white and previously characterized TEs in *Heliconius melpomene* in the RepeatMasker library 4.0.8 (Bao et al. 2015).

### Sampling of individuals

Adult male wood whites were collected across the distribution range in Sweden during June and July 2020 (Supplementary Table 4). Sex was determined *in situ* based on two sexually dimorphic characters; the presence of a black apical spot on the forewing and the white coloration of the ventral part of the antennae in males. Sampled individuals were directly preserved in ethanol and frozen at −20°C.

### DNA extraction

DNA was extracted following two different protocols. In both cases, the dissected tissue was digested overnight in Laird’s buffer and homogenized with 20*μl* of proteinase K (20mg/ml, >600 mAU/ml), followed by incubation with RNase A at 37°C for 30 minutes. DNA was extracted from thoraces using salt extraction; 300 of NaCl (5M) was added, followed by centrifugation for 15 minutes at 13,000 revolutions per minute (rpm). Three washing steps were completed with one volume of 70% ethanol and centrifuging for five minutes at maximum speed. The remaining pellet was air-dried and then resuspended in 30 *μl* of MilliQ H2O. For the abdomens, a phenol-chloroform extraction protocol was used. Two cycles of phenol:chloroform:isoamyl alcohol (25:24:1) addition and centrifugation for five minutes at 13,000 rpm were completed, plus a third cleaning cycle using only chloroform. Precipitation of DNA was achieved by adding 2x volumes isopropanol + 0.1x 3M NaAc, incubating at −18°C overnight and centrifuging for 15 minutes at 13,000 rpm. The final pellet was resuspended with 30 *μl* of MilliQ H_2_O. DNA purity was assessed with NanoDrop, and concentration measured with Qubit DNA Broad Range.

### Sequencing

To capture the genetic variation in the population in Sweden, 84 individuals from different geographic regions and the highest DNA quality were selected for analysis. Library preparation for all 84 samples using the TruSeq PCR-free kit followed by multiplexing, and sequencing on two NovaSeq 6000 S4 lanes with 2×150 bp reads, were performed at the National Genomics Infrastructure (NGI), Stockholm.

### Read trimming

Illumina sequencing adapters were trimmed by eliminating the first fifteen base pairs (bp) on each end of the raw reads with CutAdapt 1.9.1 (Martin 2011), filtered on Q-score < 30 and a minimum length of 30 bp. Read quality after cleaning was assessed with FastQC (Andrews 2010). Before filtering, an average of 4.3 million reads per sample were obtained, and 2.5% were filtered out.

### Mapping and filtering

For each individual, paired-end reads were mapped to the reference genome with bwa v0.7.17 (Li and Durbin 2009). Samtools v1.10 (Li et al. 2009) was used to select reads with paired information. MarkDuplicatesSpark as implemented in GATK v4.1.4.1 (McKenna et al. 2010) was used to eliminate duplicated regions with the –remove-sequencing-duplicates option.

### Variant calling and filtering

The tool HaplotypeCaller in GATK v4.1.4.1 (McKenna et al. 2010) was used for variant calling. Each chromosome for each individual was processed separately, and the resulting 84 files for each chromosome were grouped and converted into a VCF file with the GATK v4.1.4.1 tools Combine_gVCF and Genotype_gVCF (McKenna et al. 2010), respectively. Total variant count was obtained with the stats option in bcftools v1.10 (Li 2011). The variants were filtered to have a minimum minor allele count (MAC) of two, a per-site depth between 10 and 50, minimum per site quality of 30, and < 20% per-site missing data with vcftools 0.1.15 (Danecek et al. 2011). Additionally, all insertions and deletions were removed with the –remove-indels option. The number of remaining sites in each chromosome was counted again with the stats option in bcftools v1.10 (Li 2011). Initial variant calling resulted in a total of 51,189,479 markers along the genome (11,055,543 indels + 40,133,936 SNPs), of which 10,565,404 SNPs remained after filtering. This represents a genome-wide average of ~17.6 SNPs/kb, given the ~ 600 Mb total length of the reference genome.

### Inference of the demographic history

SMC++ (Terhorst et al. 2017) was used to infer the demographic history of each individual chromosome, using a set of six “distinguished” individuals. These six individuals were selected among the 10 with a higher average sequencing coverage for the variants after filtering, so that they constituted a good representation of the geographic distribution of the species. The per-base mutation rate was set at 2.9 x 10-9 per generation, an estimate based on mutation frequency in *H. melpomene* (Keightley et al. 2015). Known invariable regions such as centromeres must be masked before inferring the demographic history, as they can interfere with the signal. Since Lepidoptera are holocentric and lack defined centromeres, a cut-off value of 150 kb was set instead, so that any longer invariable region was considered as missing data and discarded for the demographic inference. The demographic trajectories were inferred for the last 5 million generations, as defaulted by the program.

### Recombination rate estimation

The chromosome-specific demographic trajectories were used together with the VCF files to obtain high-resolution recombination maps using pyrho (Spence and Song 2019). An algorithm implemented in the software LDpop (Kamm et al. 2016) was used to compute a table of two-loci likelihoods under the coalescent with recombination using the chromosome-specific demographic trajectories as input. The same mutation rate as before was used, together with the parameter values n = 168 (twice the number of diploid individuals), and N = 210 (25% larger than n, as recommended by the manual). A relative tolerance (`--decimate_rel_tol`) value of 0.1 was used, together with the `--approx flag`, recommended for large datasets. Different window sizes (maximum distance between SNPs) and the block penalty (determinant of the smoothness of the curve) were tested for each chromosome, and the most appropriate were selected based on the correlation between the data in the likelihood tables and simulated data at different scales (1 bp, 10 kb and 100 kb). For all chromosomes, the best block_penalty was 25, and the best suitable window_size ranged between 50 and 100. The look-up table, together with the final VCF file for each chromosome were used to infer the local recombination rate using the most appropriate parameter values and the `--fast_missing` flag. The per-base, per-generation recombination rate between each pair of SNP markers was obtained in the end. A positive association between the recombination rate and marker density was observed at a 1Mb scale (see Supplementary Information and Supplementary Figure 4 for further discussion).

### Distribution of recombination rate variation and identification of hot-spots and cold-spots

Regional recombination rate estimates were obtained in windows of two different sizes (100 kb and 1 Mb) with a custom script by calculating the weighted mean on each interval between markers, accounting for their length and recombination rate. We used simulation-based approach to establish thresholds for hot-spot identification. By performing coalescent simulation with the flat recombination landscape but taking into account previously inferred population history and genetic drift and we obtained levels of recombination rate variation, which are due to our imputation strategy. msprime 1.1.1 (Kelleher et al. 2016) was used to simulate 99 independent sequences, each 100 kb long, and the VCF files for each sequence were concatenated. The parameters for the simulations included a flat recombination landscape (7.37 cM/Mb, same as the obtained genome-wide rate) and the same mutation rate used for SMC++, together with a demographic trajectory that approximately reflects the inferred trajectories from the empirical data; exponential growth in the interval 104-106 generations BP, exponential decline 102-104 generations BP, and stable population in the last 102 generations. At each time point, *N_e_* was calculated as the average of the estimates across chromosomes. The simulated genomic sequences were analyzed in pyrho according to the steps described earlier. The resulting regional recombination rates predominantly oscillated in the range 2-8 cM/Mb, with a maximum of 20 cM/Mb, a 3-fold higher rate compared to the genome-wide average (Supplementary Figure 5). Recombination peaks occurring at boundaries between independently obtained VCF files were omitted. Therefore, simulation allowed us to establish lower bound for hot-spot threshold. For the final (more conservative) threshold we choose regions with a recombination rate higher than 25 cM/Mb, between 750 and 10,000 bp long, and showing a 10-fold increase over the regional background recombination rate (the mean rate in the focal 100 kb window and the two flanking windows, in total 300 kb). To avoid biases resulting from erroneously called variants, hot-spots that included less than four markers (i.e., three intervals) were discarded. Recombination cold-spots were identified as regions with a recombination rate 10-fold lower than the genome-wide average, and including 4 markers with no length limitations.

For the assessment of the distribution of recombination hotspots and cold-spots, we defined the subtelomeric regions as the last 500 kb on each chromosome end. The rest of the genome was considered proximal. Flanking regions to recombination hot-spots were defined as the 5 kb segments on each side of each accepted hot-spot.

### Association of recombination rate with GC content, gene density and TE classes

A multiple linear regression model was constructed to disentangle the explanatory potential of each genomic feature in the variation of recombination rate at a 1 Mb scale, using the lm function in base R (R Core Team 2021). The linear model had the recombination rate as response variable, and the explanatory variables included the GC content, the gene density, and the relative abundance of four TE classes (DNA transposons, and LINE, SINE and LTR retrotransposons). Potential associations between the regional recombination rate and the different genomic features were also analyzed with Spearman’s rank correlation tests using the corr option in base R (R Core Team 2021).

BEDTools 2.29.2 (Quinlan and Hall 2010) maskfasta option was used to select all annotated exons and TEs. Base composition for specific regions (corrected for masked positions) was obtained with BEDTools nuc option. Gene and TE densities were calculated as the proportion of a region covered by the annotated sequences of each category.

To assess potential variation in the recombination rate within specific genomic features, we also estimated the recombination rate within each TE class, different regions in protein coding genes (exons, introns and upstream regions) and intergenic sequence. Since 5’ UTR regions were not included in the annotation file, we used the 100 bp upstream of the first exon of each gene – a conservative selection to represent the 5’ UTR (Chen et al. 2011, 2014). In order to avoid biases due to differential selective pressures, only the intervals between markers from the pyrho output that were positioned completely within the feature were considered - i.e. for a given exon, if an interval between markers overlapped completely with the predicted exon, it was retained; if the overlap was only partial (including part of the exon sequence but also part of a neighboring intron or UTR), it was discarded. This overlap was checked with BEDTools 2.29.2 (Quinlan and Hall 2010) intersect option using the flag -f 1.

### Permutation test to assess TE density in the cold- and hot-spots

To assess the TE density in the cold and hotspot regions we ran a permutation test in R v4.2.1 (R Core Team, 2013), by resampling 10,000 means of windows reflecting the count and average length of the outlier windows and assessing where in this distribution of resampled means the value of the hot- and cold-spot windows appears. We then performed a two-sided test by assessing the number of means that exceeded the original difference.

Gene Set Enrichment Analysis (GSEA) for the genes present in the cold- and hot-spot regions We assessed enrichment of functional categories in the cold and hot spot regions using topGO v2.44.0 (Alexa and Rahnenfuhrer 2021) in R v4.2.1 (R Core Team, 2013). We used the annotated gene set with gene ontology (GO) terms associated to molecular function. To assess significance, we used the Fisher’s exact test and the default algorithm (“weight01”) accounting for the hierarchical structure of the GO-terms (Alexa et al. 2006). We adjusted the p-values with Benjamini-Hochberg’s method of multiple test correction (p.adjust(x, method = “fdr”)). We used HOMER v4.11 (Heinz et al. 2010) to assess motif enrichment in the cold and hot spot windows.

## Data access

All raw data generated in this study has been submitted to the European Nucleotide Archive (ENA; https://www.ebi.ac.uk/ena/browser/home) under accession number PRJEB56690. Command lines are available on the group’s GitHub page (https://github.com/EBC-butterfly-genomics-team).

## Acknowledgements

This work was supported by a research grant from the Swedish Research Council (Vetenskapsrådet Grant ID: 2019-04791) to N.B. The authors acknowledge support from the National Genomics Infrastructure in Stockholm funded by Science for Life Laboratory, the Knut and Alice Wallenberg Foundation and the Swedish Research Council, and SNIC/Uppsala Multidisciplinary Center for Advanced Computational Science for assistance with massively parallel sequencing and access to the UPPMAX computational infrastructure. This work was also supported by NBIS/SciLifeLab long-term bioinformatics support (WABI). R.V. was supported by grant PID2019-107078GB-I00, funded by MCIN/AEI/10.13039/ 501100011033.

