## Supplemental Material for "The fine-scale recombination rate variation and associations with genomic features in a butterfly"

### Supplementary Material: Table of Contents

**SUPPLEMENTARY INFORMATION** **1**

**SUPPLEMENTARY INFORMATION**

*Recent demographic trends are the consequence of Post-Pleistocene population dynamics*

The inferred demographic trajectories were consistent across chromosomes. The historical fluctuations in $N_e$ were in agreement with previous analyses, with a gradual increase until approximately 10,000 generations ago (Talla *et al.* 2019). By using a method (SMC++) that can accomodate unphased data from many individuals, we could infer trajectories in the recent past (Terhorst *et al.* 2017). We found a considerable decrease in $N_e$ in the last 10,000 generations, a signal that Talla *et al.* (2019) could not pick up with PSMC (Li and Durbin 2011), which is not designed for inferences of $N_e$ in the recent past as it uses genomic information from a single individual (Mather *et al.* 2020). The perceived decline in $N_e$ during the last 10,000 generations is consistent with recurrent founder effects during recolonization of Scandinavia after the latest glaciation period and is compatible with the hypothesis of secondary contact and recurrent back-crossing underlying the observed chromosome number cline observed across Eurasia (Lukhtanov *et al.* 2018; Talla *et al.* 2019).

An important consideration regarding the time estimates of the demographic changes is that the assumption of univoltinity may be a simplification of the true scenario. Currently, wood whites is univoltine in the northern parts of the distribution range (Friberg *et al.* 2008), but the number of annual generations increases further south (Jeffcoate and Joy 2011, Dantart 2003). Historical changes in voltinity (e.g. Margus and Lindström 2020, Powell 1989) could affect the scale of inferences on population size evolution. However, the timing of the ice sheet retreat in Scandinavia is estimated at about 11,000 years BP (Günther *et al.* 2018), which matches the beginning of the observed decrease in $N_e$. This indicates that the decrease in $N_e$ is most likely a consequence of founder effects during the recolonization of Scandinavia from glacial refugia.

*Local recombination rate is associated with marker density*

We found that marker density was positively correlated with the estimated recombination rate at a 1 Mb scale, when omitting regions at the very end of chromosomes (Supplementary Figure 4). This association has also been observed in other systems (Shen *et al.* 2018), due to the decrease in genetic variation caused by deleterious sweeps (Maynard Smith and Haigh 1974, Jacobs *et al.* 2016). In these regions, the estimated recombination rate can be lower than the real value (Kim and Nielsen 2004).

#### SUPPLEMENTARY TABLES

**Supplementary Table 1. The recombination rate decreases in most subtelomeric regions.** *p-values* for the Wilcoxon tests, with values in with an asterisk indicating a non-significant decrease in the estimated recombination rate compared to the central positions of the chromosome.

| Chromosome | ‘Left’ subtelomeric region | ‘Right’ subtelomeric region |
|---|---|---|
| Z1 | <0.001 | 0.014 |
| 2 | <0.001 | 0.004 |
| 3 | <0.001 | 0.025 |
| 4 | 0.001 | 0.001 |
| 5 | 0.085* | 0.008 |
| Z2 | 0.002 | 0.731* |
| 7 | 0.008 | 0.007 |
| 8 | 0.023 | 0.003 |
| 9 | 0.017 | 0.033 |
| 10 | 0.762* | 0.001 |
| 11 | 0.524* | 0.001 |
| 12 | 0.001 | <0.001 |
| 13 | 0.001 | 0.002 |
| 14 | <0.001 | <0.001 |
| 15 | 0.0256 | <0.001 |
| 16 | 0.950* | 0.014 |
| 17 | 0.002 | <0.001 |
| 18 | 0.015 | 0.009 |
| Z3 | 0.017 | 0.005 |
| 20 | 0.005 | 0.012 |
| 21 | 0.002 | 0.004 |
| 22 | 0.009 | 0.003 |
| 23 | 0.002 | <0.001 |
| 24 | 0.001 | 0.002 |

| | | |
|---|---|---|
| 25 | 0.001 | 0.098* |
| 26 | <0.001 | 0.001 |
| 27 | 0.005 | 0.098* |
| 28 | <0.001 | 0.003 |
| 29 | 0.246* | 0.004 |

**Supplementary Table 2. GSEA output for the 50 most significant GO terms in the hot-spots.** Shown for each GO term is the name, total number of genes belonging to this GOterm, expected, and observed amount in the hot-spots, p-value and adjusted p-value for each GOterm ID.

| **GO.ID** | **Term** | **#Annot.** | **#Sign.** | **#Exp.** | **Fisher** | **p_adj** |
|---|---|---|---|---|---|---|
| GO:0005219 | ryanodine-sensitive calcium-release channel activity | 3 | 3 | 0.18 | 2.20E-04 | 0.37 |
| GO:0043924 | suramin binding | 3 | 3 | 0.18 | 2.20E-04 | 0.37 |
| GO:0005509 | calcium ion binding | 159 | 21 | 9.64 | 5.70E-04 | 0.57 |
| GO:0050254 | rhodopsin kinase activity | 4 | 3 | 0.24 | 8.50E-04 | 0.57 |
| GO:0047696 | beta-adrenergic receptor kinase activity | 4 | 3 | 0.24 | 8.50E-04 | 0.57 |
| GO:0005524 | ATP binding | 204 | 24 | 12.37 | 1.27E-03 | 0.65 |
| GO:0048763 | calcium-induced calcium release activity | 5 | 3 | 0.3 | 2.02E-03 | 0.65 |
| GO:0000400 | four-way junction DNA binding | 11 | 4 | 0.67 | 3.13E-03 | 0.65 |
| GO:0034237 | protein kinase A regulatory subunit binding | 11 | 4 | 0.67 | 3.13E-03 | 0.65 |
| GO:0031850 | delta-type opioid receptor binding | 2 | 2 | 0.12 | 3.67E-03 | 0.65 |
| GO:0031851 | kappa-type opioid receptor binding | 2 | 2 | 0.12 | 3.67E-03 | 0.65 |
| GO:0046702 | galactoside 6-L-fucosyltransferase activity | 2 | 2 | 0.12 | 3.67E-03 | 0.65 |
| GO:0050614 | delta24-sterol reductase activity | 2 | 2 | 0.12 | 3.67E-03 | 0.65 |
| GO:0000246 | delta24(24-1) sterol reductase activity | 2 | 2 | 0.12 | 3.67E-03 | 0.65 |
| GO:0046935 | 1-phosphatidylinositol-3-kinase regulator activity | 2 | 2 | 0.12 | 3.67E-03 | 0.65 |
| GO:0008732 | L-allo-threonine aldolase activity | 2 | 2 | 0.12 | 3.67E-03 | 0.65 |
| GO:0031755 | Edg-2 lysophosphatidic acid receptor binding | 2 | 2 | 0.12 | 3.67E-03 | 0.65 |
| GO:0008424 | glycoprotein 6-alpha-L-fucosyltransferase activity | 2 | 2 | 0.12 | 3.67E-03 | 0.65 |
| GO:0005161 | platelet-derived growth factor receptor binding | 6 | 3 | 0.36 | 3.86E-03 | 0.65 |
| GO:0004703 | G protein-coupled receptor kinase activity | 6 | 3 | 0.36 | 3.86E-03 | 0.65 |

| | | | | | | |
|---|---|---|---|---|---|---|
| GO:0050839 | cell adhesion molecule binding | 120 | 13 | 7.28 | 5.10E-03 | 0.82 |
| GO:0050062 | long-chain-fatty-acyl-CoA reductase activity | 7 | 3 | 0.42 | 6.45E-03 | 0.90 |
| GO:0003993 | acid phosphatase activity | 7 | 3 | 0.42 | 6.45E-03 | 0.90 |
| GO:0034236 | protein kinase A catalytic subunit binding | 7 | 3 | 0.42 | 6.45E-03 | 0.90 |
| GO:0047372 | acylglycerol lipase activity | 8 | 3 | 0.49 | 9.86E-03 | 1.00 |
| GO:0042605 | peptide antigen binding | 3 | 2 | 0.18 | 1.06E-02 | 1.00 |
| GO:0000403 | Y-form DNA binding | 3 | 2 | 0.18 | 1.06E-02 | 1.00 |
| GO:0031694 | alpha-2A adrenergic receptor binding | 3 | 2 | 0.18 | 1.06E-02 | 1.00 |
| GO:0005245 | voltage-gated calcium channel activity | 18 | 5 | 1.09 | 1.06E-02 | 1.00 |
| GO:0001530 | lipopolysaccharide binding | 15 | 4 | 0.91 | 1.07E-02 | 1.00 |
| GO:0005085 | guanyl-nucleotide exchange factor activity | 119 | 14 | 7.22 | 1.25E-02 | 1.00 |
| GO:0043621 | protein self-association | 54 | 8 | 3.28 | 1.52E-02 | 1.00 |
| GO:0097110 | scaffold protein binding | 45 | 7 | 2.73 | 1.76E-02 | 1.00 |
| GO:0001968 | fibronectin binding | 10 | 3 | 0.61 | 1.93E-02 | 1.00 |
| GO:0031492 | nucleosomal DNA binding | 4 | 2 | 0.24 | 2.03E-02 | 1.00 |
| GO:0004663 | Rab geranylgeranyltransferase activity | 4 | 2 | 0.24 | 2.03E-02 | 1.00 |
| GO:0009374 | biotin binding | 4 | 2 | 0.24 | 2.03E-02 | 1.00 |
| GO:0017108 | 5'-flap endonuclease activity | 4 | 2 | 0.24 | 2.03E-02 | 1.00 |
| GO:0008349 | MAP kinase kinase kinase kinase activity | 4 | 2 | 0.24 | 2.03E-02 | 1.00 |
| GO:0003700 | DNA-binding transcription factor activity | 474 | 34 | 28.75 | 2.76E-02 | 1.00 |
| GO:0070412 | R-SMAD binding | 20 | 4 | 1.21 | 2.98E-02 | 1.00 |
| GO:0034041 | ABC-type sterol transporter activity | 12 | 3 | 0.73 | 3.23E-02 | 1.00 |
| GO:0030619 | U1 snRNA binding | 5 | 2 | 0.3 | 3.25E-02 | 1.00 |
| GO:0004704 | NF-kappaB-inducing kinase activity | 5 | 2 | 0.3 | 3.25E-02 | 1.00 |
| GO:0019209 | kinase activator activity | 51 | 4 | 3.09 | 3.26E-02 | 1.00 |
| GO:0001872 | (1->3)-beta-D-glucan binding | 6 | 2 | 0.36 | 4.68E-02 | 1.00 |

| | | | | | | |
|---|---|---|---|---|---|---|
| GO:0005242 | inward rectifier potassium channel activity | 6 | 2 | 0.36 | 4.68E-02 | 1.00 |
| GO:0043422 | protein kinase B binding | 6 | 2 | 0.36 | 4.68E-02 | 1.00 |
| GO:0043199 | sulfate binding | 6 | 2 | 0.36 | 4.68E-02 | 1.00 |
| GO:0080019 | fatty-acyl-CoA reductase (alcohol-forming) activity | 14 | 3 | 0.85 | 4.89E-02 | 1.00 |

**Supplementary Table 3. GSEA output for the 50 most significant GO terms in the cold-spots.** Shown for each GO term is the name, total number of genes belonging to this GOterm. expected and observed amount in the cold spots, p-value and adjusted p-value for each GOterm ID. Significantly enriched GOterms are highlighted in bold.

| GO.ID | Term | Annot. | Sign | Expected | weight01Fisher | p_adj |
|---|---|---|---|---|---|---|
| **GO:0004311** | **farnesyltranstransferase activity** | **21** | **13** | **1.50** | **9.70E-09** | **0.00** |
| **GO:0004337** | **geranyltranstransferase activity** | **19** | **11** | **1.36** | **1.00E-08** | **0.00** |
| **GO:0004161** | **dimethylallyltranstransferase activity** | **19** | **11** | **1.36** | **1.00E-08** | **0.00** |
| GO:0004467 | long-chain fatty acid-CoA ligase activity | 17 | 7 | 1.22 | 9.50E-05 | 0.08 |
| GO:0004370 | glycerol kinase activity | 6 | 4 | 0.43 | 3.50E-04 | 0.20 |
| GO:0046870 | cadmium ion binding | 3 | 3 | 0.21 | 3.60E-04 | 0.20 |
| GO:0004521 | endoribonuclease activity | 43 | 8 | 3.08 | 1.10E-03 | 0.37 |
| GO:0030899 | calcium-dependent ATPase activity | 4 | 3 | 0.29 | 1.38E-03 | 0.37 |
| GO:0004517 | nitric-oxide synthase activity | 4 | 3 | 0.29 | 1.38E-03 | 0.37 |
| GO:0034617 | tetrahydrobiopterin binding | 4 | 3 | 0.29 | 1.38E-03 | 0.37 |
| GO:0034618 | arginine binding | 4 | 3 | 0.29 | 1.38E-03 | 0.37 |
| GO:0004169 | dolichyl-phosphate-mannose-protein mannosyltransferase activity | 4 | 3 | 0.29 | 1.38E-03 | 0.37 |
| GO:0017080 | sodium channel regulator activity | 8 | 4 | 0.57 | 1.44E-03 | 0.37 |
| GO:0008603 | cAMP-dependent protein kinase regulator activity | 5 | 3 | 0.36 | 3.27E-03 | 0.67 |
| GO:1904047 | S-adenosyl-L-methionine binding | 5 | 3 | 0.36 | 3.27E-03 | 0.67 |
| GO:0008449 | N-acetylglucosamine-6-sulfatase activity | 5 | 3 | 0.36 | 3.27E-03 | 0.67 |
| GO:0005351 | carbohydrate:proton symporter activity | 29 | 7 | 2.08 | 3.61E-03 | 0.67 |
| GO:0005355 | glucose transmembrane transporter activity | 29 | 7 | 2.08 | 3.61E-03 | 0.67 |
| GO:0003857 | 3-hydroxyacyl-CoA dehydrogenase activity | 10 | 4 | 0.72 | 3.85E-03 | 0.68 |
| GO:0016275 | [cytochrome c]-arginine N-methyltransferase activity | 2 | 2 | 0.14 | 5.12E-03 | 0.69 |

| | | | | | | |
|---|---|---|---|---|---|---|
| GO:0042071 | leucokinin receptor activity | 2 | 2 | 0.14 | 5.12E-03 | 0.69 |
| GO:0005152 | interleukin-1 receptor antagonist activity | 2 | 2 | 0.14 | 5.12E-03 | 0.69 |
| GO:0008495 | protoheme IX farnesyltransferase activity | 2 | 2 | 0.14 | 5.12E-03 | 0.69 |
| GO:0030519 | snoRNP binding | 2 | 2 | 0.14 | 5.12E-03 | 0.69 |
| GO:0070224 | sulfide:quinone oxidoreductase activity | 2 | 2 | 0.14 | 5.12E-03 | 0.69 |
| GO:0043199 | sulfate binding | 6 | 3 | 0.43 | 6.19E-03 | 0.77 |
| GO:0003958 | NADPH-hemoprotein reductase activity | 6 | 3 | 0.43 | 6.19E-03 | 0.77 |
| GO:0003887 | DNA-directed DNA polymerase activity | 19 | 5 | 1.36 | 9.24E-03 | 1.00 |
| GO:0036122 | BMP binding | 7 | 3 | 0.50 | 1.03E-02 | 1.00 |
| GO:0016887 | ATP hydrolysis activity | 343 | 36 | 24.56 | 1.22E-02 | 1.00 |
| GO:0020037 | heme binding | 36 | 7 | 2.58 | 1.25E-02 | 1.00 |
| GO:0004222 | metalloendopeptidase activity | 45 | 8 | 3.22 | 1.33E-02 | 1.00 |
| GO:0016618 | hydroxypyruvate reductase activity | 3 | 2 | 0.21 | 1.46E-02 | 1.00 |
| GO:0030267 | glyoxylate reductase (NADP+) activity | 3 | 2 | 0.21 | 1.46E-02 | 1.00 |
| GO:0050660 | flavin adenine dinucleotide binding | 48 | 8 | 3.44 | 1.50E-02 | 1.00 |
| GO:0008409 | 5'-3' exonuclease activity | 12 | 4 | 0.86 | 2.20E-02 | 1.00 |
| GO:0015036 | disulfide oxidoreductase activity | 32 | 4 | 2.29 | 2.22E-02 | 1.00 |
| GO:0004518 | nuclease activity | 112 | 15 | 8.02 | 2.73E-02 | 1.00 |
| GO:0036185 | 13-lipoxin reductase activity | 4 | 2 | 0.29 | 2.79E-02 | 1.00 |
| GO:0035175 | histone kinase activity (H3-S10 specific) | 4 | 2 | 0.29 | 2.79E-02 | 1.00 |
| GO:0051400 | BH domain binding | 4 | 2 | 0.29 | 2.79E-02 | 1.00 |
| GO:0097257 | leukotriene B4 12-hydroxy dehydrogenase activity | 4 | 2 | 0.29 | 2.79E-02 | 1.00 |
| GO:0004321 | fatty-acyl-CoA synthase activity | 4 | 2 | 0.29 | 2.79E-02 | 1.00 |
| GO:0097371 | MDM2/MDM4 family protein binding | 4 | 2 | 0.29 | 2.79E-02 | 1.00 |
| GO:0044020 | histone methyltransferase activity (H4-R3 specific) | 4 | 2 | 0.29 | 2.79E-02 | 1.00 |
| GO:0032027 | myosin light chain binding | 10 | 3 | 0.72 | 3.00E-02 | 1.00 |

| | | | | | | |
|---|---|---|---|---|---|---|
| GO:0010181 | FMN binding | 10 | 3 | 0.72 | 3.00E-02 | 1.00 |
| GO:0016303 | 1-phosphatidylinositol-3-kinase activity | 11 | 3 | 0.79 | 3.91E-02 | 1.00 |
| GO:0003785 | actin monomer binding | 11 | 3 | 0.79 | 3.91E-02 | 1.00 |
| GO:0032450 | maltose alpha-glucosidase activity | 5 | 2 | 0.36 | 4.43E-02 | 1.00 |

**Supplementary Table 4. Sampling coordinates of the sequenced individuals.**

| Location | Latitude | Longitude | # Individuals |
|---|---|---|---|
| Fiby | N 59° 54' 08.01" | E 17° 21' 31.76" | 2 |
| Lumpen | N 59° 57' 00.53" | E 17° 18' 30.05” | 2 |
| Stora Trillbo | N 59° 54' 49.79" | E 17° 23' 28.16" | 2 |
| Siggefora | N 59° 58' 46.24" | E 17° 07' 20.56" | 2 |
| Östfora | N 59° 59' 22.41" | E 17° 10' 47.80" | 1 |
| Vattholma | N 60° 01' 31.52" | E 17° 45' 58.96" | 3 |
| Lafsen | N 60° 01' 36.45" | E 17° 48' 08.62" | 2 |
| Södra Lafsen | N 60° 01' 00.62" | E 17° 49' 01.83" | 2 |
| Översjön Jarva | N 59° 27' 23.32" | E 17° 51' 02.12" | 1 |
| Fjärilsstigen | N 59° 50' 13.58" | E 17° 32' 46.84" | 2 |
| Västmanland | N 59° 42' 55.55" | E 16° 05' 35.45" | 4 |
| Närke | N 59° 19' 25.25" | E 14° 45' 19.11" | 7 |
| Värmland | N 59° 39' 56.24" | E 13° 32' 57.27" | 9 |
| Dalarna | N 60° 08' 01.02" | E 15° 15' 47.47" | 8 |
| Hälsingland | N 61° 22' 30.03" | E 16° 16' 37.23" | 4 |
| Roxen | N 58° 17' 08.50" | E 14° 55' 35.60" | 1 |
| Forserum | N 57° 43' 29.00" | E 14° 30' 45.10" | 1 |
| Flögen | N 57° 25' 58.20" | E 14° 59' 19.80" | 5 |
| Landsbro | N 57° 22' 08.50" | E 14° 54' 13.80" | 1 |
| Rottnen | N 56° 42' 37.40" | E 15° 05' 45.90" | 3 |
| Norrbo | N 60° 16' 31.9" | E 14° 53' 58.1" | 3 |
| Söraker | N 62° 29' 17.51" | E 17° 36' 35.72" | 6 |
| Varmvattnet | N 64° 04' 40.94" | E 20° 03' 20.52" | 4 |
| Malå | N 65° 09' 21.81" | E 18° 46' 58.99" | 1 |
| Föllinge | N 63° 34' 12.28" | E 14° 43' 04.12" | 8 |

#### SUPPLEMENTARY FIGURES

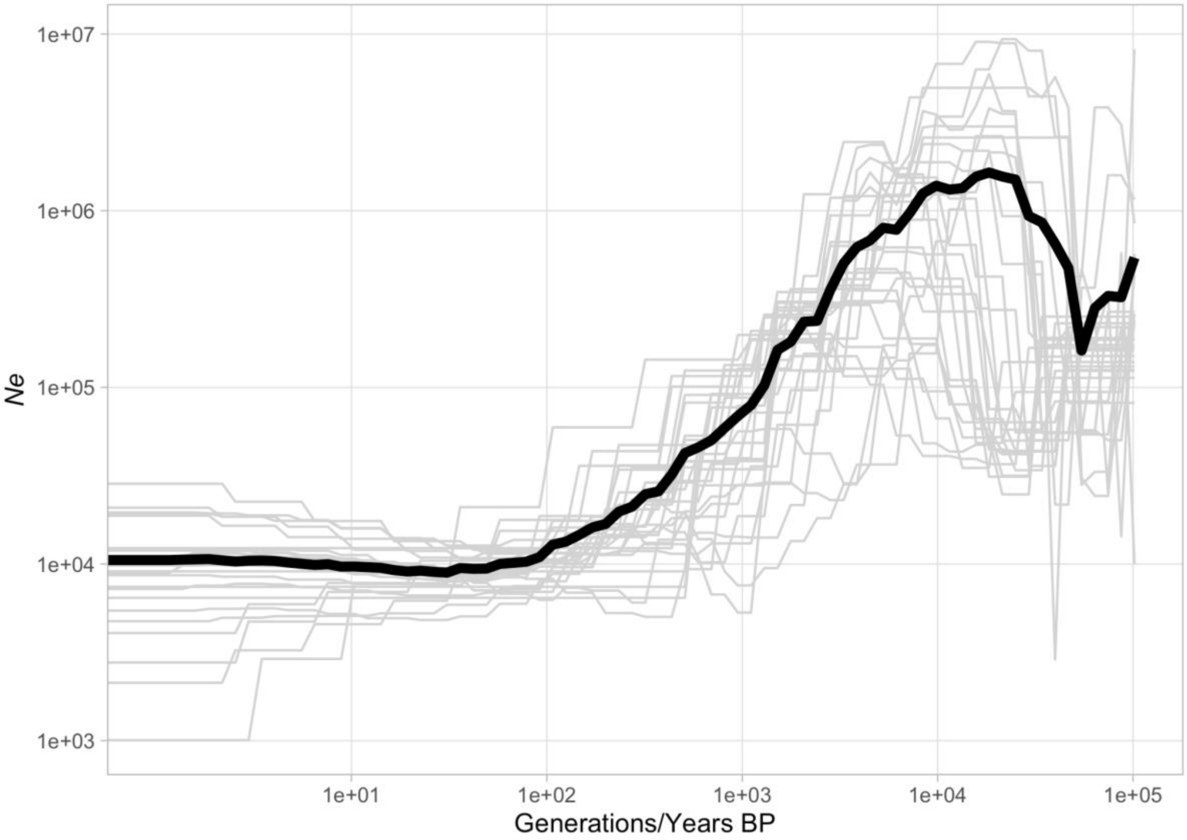

**Supplementary Figure 1. Demographic history inferred for the different chromosomes in the Swedish population of wood white.** The last 100,000 generations of the demography are shown, as all variation in concentrated in that range. This is equivalent to the same number of years if we assume one annual generation in Scandinavia. The black line indicates the mean value for $N_e$ at different time points.

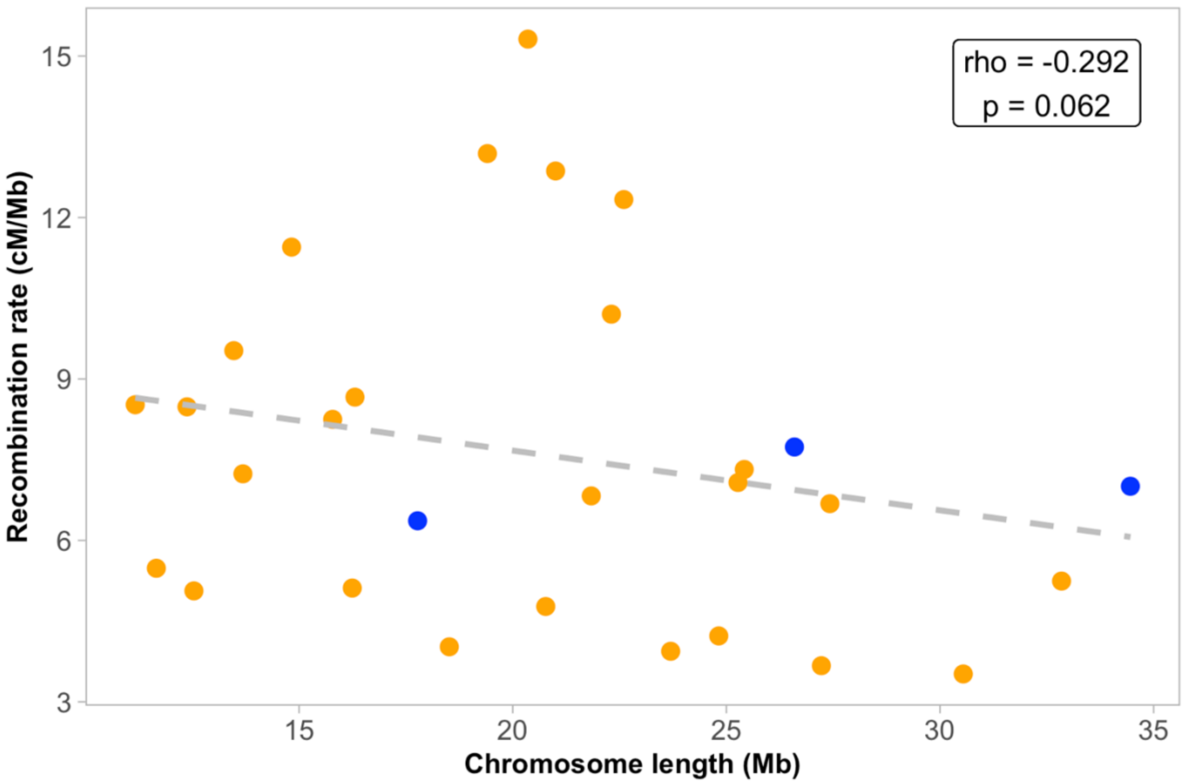

**Supplementary Figure 2. The association between chromosome length and recombination rate in the wood white.** Each dot represents a chromosome (orange = autosomes, blue = Z-chromosomes).

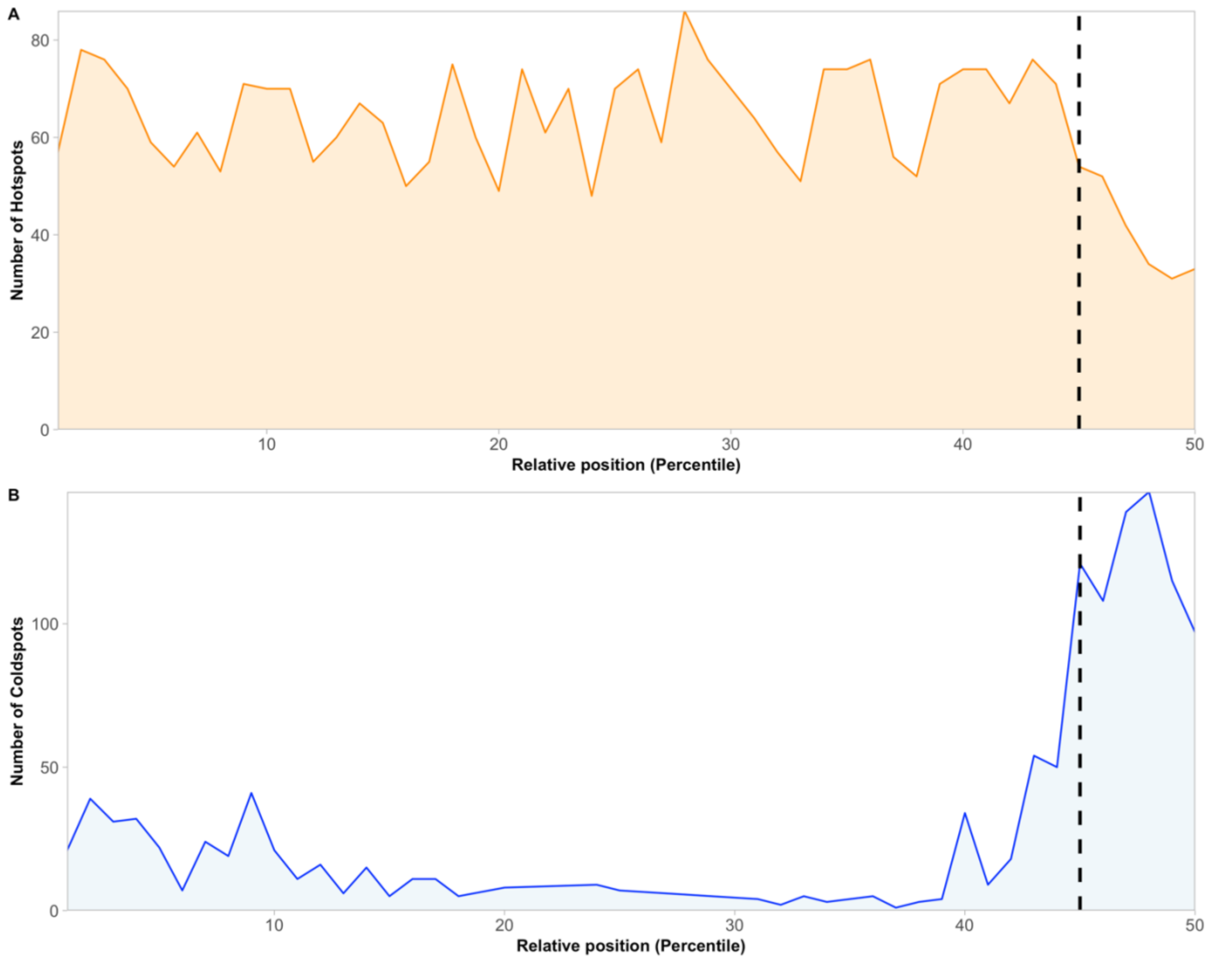

**Supplementary Figure 3. Relative position distribution of recombination hot-spots and cold-spots**. Distribution of recombination hot-spots **(A)** and cold-spots **(B)** along the different chromosomes, represented as the count in the percentiles of relative position with respect to the center of the chromosome.

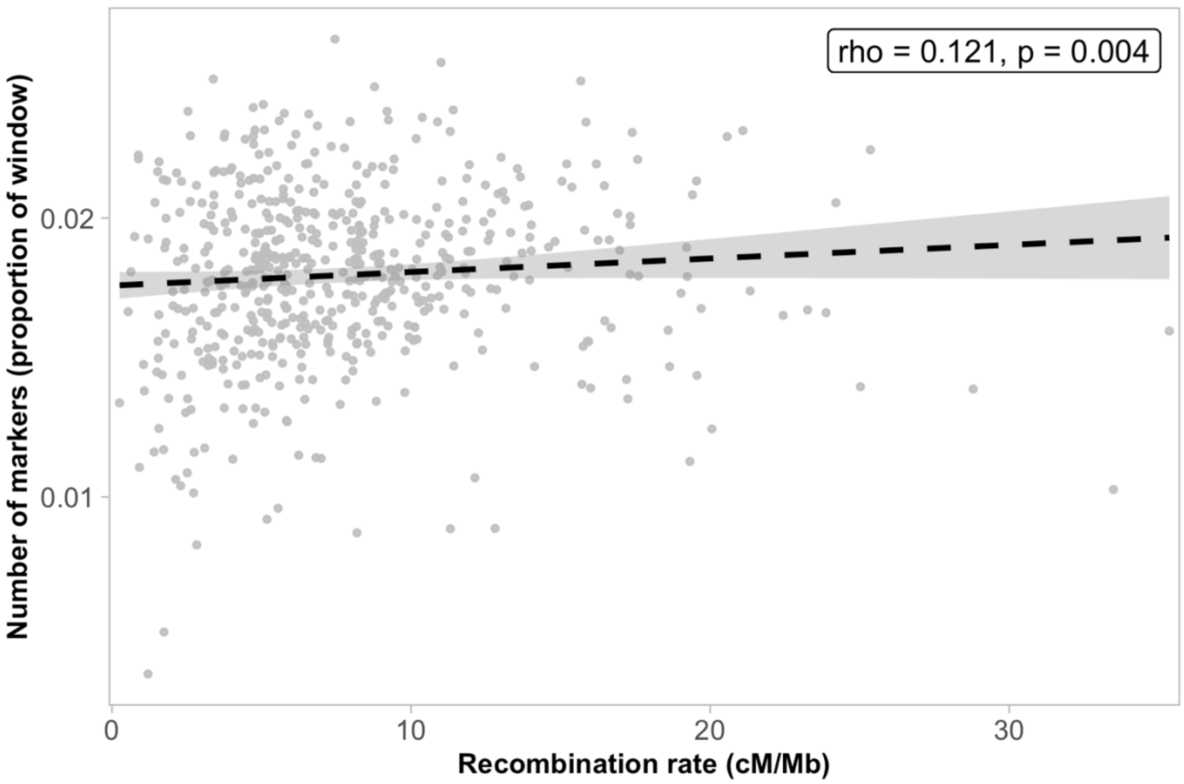

**Supplementary Figure 4. Positive association between the local recombination rate and the marker density**. The 1Mb window with the highest recombination rate (located in chromosome 17, see Figure S3) was discarded for the analysis, as well as the first and last window of each chromosome, as they included the telomeric regions.

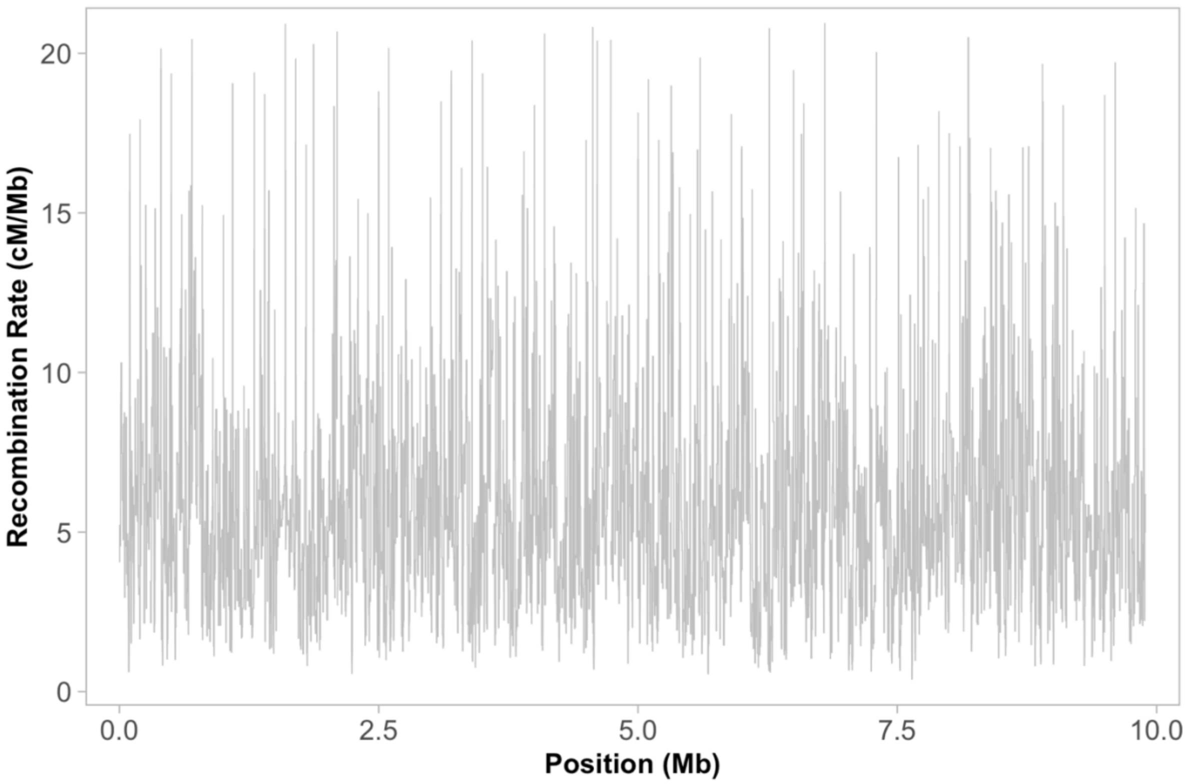

**Supplementary Figure 5.** ***pyrho*** **output for simulated data with** ***msprime***. *pyrho* was used to infer the local recombination rate variation in a single file obtained from concatenating 99 independently simulated VCF files with *msprime* 1.1.1, for a total of 9.9 Mb long sequence. Based on these results, the thresholds for defining and characterizing the recombination hot-spots were decided.
